# The White Sea littoral oligochaete *Lumbricillus* sp. as a model for annelid development studies

**DOI:** 10.1101/2024.11.19.624369

**Authors:** E.P. Matveicheva, T.V. Neretina, I.A. Ekimova, V.V. Konduktorova, M.L. Semenova, D.A. Nikishin

## Abstract

Oligochaetes play a significant role in aquatic ecosystems, and studying their development can contribute to a deeper understanding of the evolutionary features of embryonic development in the phylum Annelida. In this paper, we initially explored a littoral oligochaete species from the White Sea, which is abundant in the vicinity of the White Sea Biological Station MSU and reproduces in the early summer. A molecular phylogenetic analysis utilizing COI and 18S markers confirmed that the species under investigation belongs to the genus *Lumbricillus*, family Enchytridae. The early stages of embryonic development were described in detail, and the timing of the onset of key stages was established. The specifics of the cellular patterns observed in the early embryo were elucidated using the fluorescence staining and confocal microscopy. The embryonic development of *Lumbricillus* sp. exhibits characteristics that are consistent with the typical development of oligochaetes. Additionally, it displays specific characteristics that are typical of representatives of the Enchytriidae family, including equal size of the 2nd and 2D blastomeres and a small size of the 4D blastomere in comparison to the large 4D blastomere. The combination of synchrony and low developmental variability, coupled with the accessibility of the object, ease of maintenance and manipulation, makes *Lumbricillus* sp. a highly promising object for the study of embryonic development in Annelides.

## INTRODUCTION

Clitellata, combining leeches and oligochaetes, are considered to be a highly evolved monophyletic group of annelids of aquatic origin with adaptations to freshwater and terrestrial habitats (Rousset et al., 2008). Oligochaetes occupy a prominent position in biology and ecology, in part because they have been used to assess the ecological quality of aquatic ecosystems. However, despite their importance, ecological and biological studies on marine oligochaetes are scarce (Giere, 2006). Several biotic indices based on the analysis of oligochaete communities have been proposed for the environmental assessment of water quality. For example, the embryotoxicity test using *Enchytraeus crypticus* has been proposed (Gonçalves et al., 2015). Adaptations to diverse ecological niches often entail significant modifications in the processes of embryogenesis, which can manifest through variations such as the formation of protective shells or altered developmental pathways. In addition, a representative set of model objects is critically needed to fully study the developmental evolution of any group. Therefore, we put forth the suggestion that a littoral oligochaete species serve as the subject of study, as it is both convenient and accessible, and represents a notable addition to the number of model study objects within the Clitellata group.

The early embryonic development of the majority of oligochaetes follows a general pattern, that is characterised by a number of notable features (Dohle, 1999). As is the case with other annelids, the embryogenesis of oligochaetes is distinguished by spiral cleavage and a multitude of characteristics associated with determinative development. These include unequal cleavage and a high importance of quadrant D in development, reflecting the early establishment of the axial organisation of the embryo. The large amount of yolk in the oocytes and the pronounced difference in size between the animal and vegetal blastomeres are also characteristic features of most oligochaetes. Finally, a striking feature of the embryonic development of oligochaetes and leeches is the formation of the pair of mesoteloblasts and the four pairs of ectoteloblasts, which form two germ bands that interlock on the ventral side during the final stage of gastrulation, which proceeds by epiboly (Dohle, 1999).

This study aims to investigate the embryonic development of various Oligochaeta species, with focus on enhancing our understanding of the evolutionary traits and developmental mechanisms within the Annelida phylum. We propose the littoral oligochaete species, which is common in the WSBS MSU region and exhibits breeding activity during the early summer months, as a promising biological target for study. Our objectives include determining the phylogenetic relationships of this species and characterizing its early embryonic development through the application of light microscopy and laser scanning confocal microscopy techniques. This integrated approach has yielded new data that may potentially provide valuable contributions to the field of evolutionary and developmental biology with regard to annelids.

## MATERIALS AND METHODS

### Sample collection

Samples were collected in 2022-24 at the White Sea Biological Station of Moscow State University (WSBS, 66° 34′ N, 33° 08′ E 66.552941, 33.103014). Adult worms and cocoons with eggs of a distinctive bright orange hue are prevalent in the lower intertidal zone of the Yeremeyevsky rapids, where they are found in considerable numbers within the bases of brown algae thalloms. The specimens were collected at low tide and subsequently extracted from the substrate surface using dissecting needles. In order to investigate the process of embryonic development, the cocoons were sorted into developmental stages and maintained in filtered seawater at a constant temperature of 10°C. The study utilized a total of approximately 50 adult worms and over 350 cocoons with embryos at varying stages of development.

### Fluorescent staining and microscopy

For the purposes of fixation and staining, the cocoons were carefully opened with the use of two insulin syringe 29G needles, and the embryos were extracted into filtered seawater that had been supplemented with 5% BSA and 10% sucrose. This allowed to preserve the morphology of the embryos. The embryos of different stages were fixed with 4% paraformaldehyde (PFA) in phosphate-buffered saline (PBS, 0.01M) at 0 °C overnight. After washes and permeabilisation in 0.1% Triton X-100 (PBST), the embryos were stained by CytoPainter Phalloidin-iFluor 488 (1:200, Abcam, UK), Propidium iodide (500 nM; Merck KGaA, Germany) or DAPI (1 μg/ml; Merck KGaA, Germany) for one hour and then washed four times in PBS. The embryos were embedded in 80% glycerol on PBS solution and analyzed using a laser scanning confocal microscope Nikon A1.

### DNA sequencing and analysis

Samples of adult worms (N=5) and cocoons (N=10) containing developing embryos were used for DNA extraction and subsequent sequencing. The Promega Wizard SV Genomic DNA Purification Kit (Promega Corporation, Madison, USA) was used for tissue lysis and DNA purification following the respective protocol. Polymerase chain reactions (PCR) of nuclear 18S rRNA and mitochondrial cytochrome c oxidase subunit I (COI) were accomplished with the standard primers (PCR conditions and primers are given in Table S1). All loci were amplified using the Encyclo PCR kit (Evrogen Joint Stock Company, Moscow, Russia). Amplification products were purified using the Promega PCR Purification Kit (Promega). Due to low amplification rate of two 18S products only the third fragment amplified from 18Sa2.0-18S9R primer pair, was used for sequencing. Sequencing was performed with a BigDye Terminator v3.1 sequencing kit (Applied Biosystems), same primers as for PCR were used. Samples were purified by ethanol precipitation, were re-suspended in 12 μl formamide (Applied Biosystems) and electrophoresed in an ABI Prism 3500 Genetic Analyser (Applied Biosystems).

All raw reads for each gene were assembled and checked for ambiguities and low-quality data in Geneious R10 (Kearse et al., 2012). Edited sequences were verified for contamination using the BLAST-n algorithm run over the GenBank nr/nt database (Altschul et al., 1990). For the phylogenetic analysis a previously published dataset for the genus *Lumbricillus* (Klinth et al., 2017) was used to place the analyzed species in the phylogenetic framework (Table S2). Sequences were aligned with the MUSCLE (Edgar, 2004) algorithm in MEGA7 (Kumar et al., 2016). Final alignments comprised 629 bp for COI and 634 bp for 18S. The best-fit nucleotide evolution models were estimated in MEGA7, in both cases the GTR+G+I model was selected. The Maximum likelihood phylogenetic analysis was performed in MEGA7 with 1000 pseudoreplicates. Final phylogenetic tree images were rendered in FigTree 1.4.0 and further visually modified in Adobe Illustrator CS2015.

## RESULTS AND DISCUSSION

In this paper, we present the first description of the early developmental stages of the marine littoral oligochaete, which occurs en masse on the Karelian coast of the White Sea in the vicinity of the WSBS MSU. Pale orange or pinkish worms up to 1.2 cm long (Fig. 1A) occur in large numbers at the base of the brown algal thalli from late May until at least mid-July. In places where adult worms congregate, large numbers of cocoons containing eggs of a characteristic yellowish-orange color are observed. All development up to hatching of the juvenile oligochaetes takes place inside these cocoons, which are attached to the substrate surface (Fig. 1B-G). Cocoons are rounded, slightly ellipsoidal, with thickenings at both ends. The average cocoon length is 1.12 mm and width 0.92 mm. The cocoon walls are dense, translucent, brownish, with longitudinal grooves. Cocoons of older stages are less brightly colored and more rounded. Each cocoon contains 4 to 11 eggs surrounded by a transparent, viscous fluid (Fig. 1B). Mature eggs, measuring 0.26-0.3 mm in diameter, are encased in a closely apposed, thin vitelline membrane. The eggs are opaque, orange-yellow in color and contain a substantial quantity of yolk distributed uniformly throughout the egg’s volume. A minor region at the animal pole, surrounding nuclear material, is devoid of the yolk (Fig. 3A).

**Figure 1.**
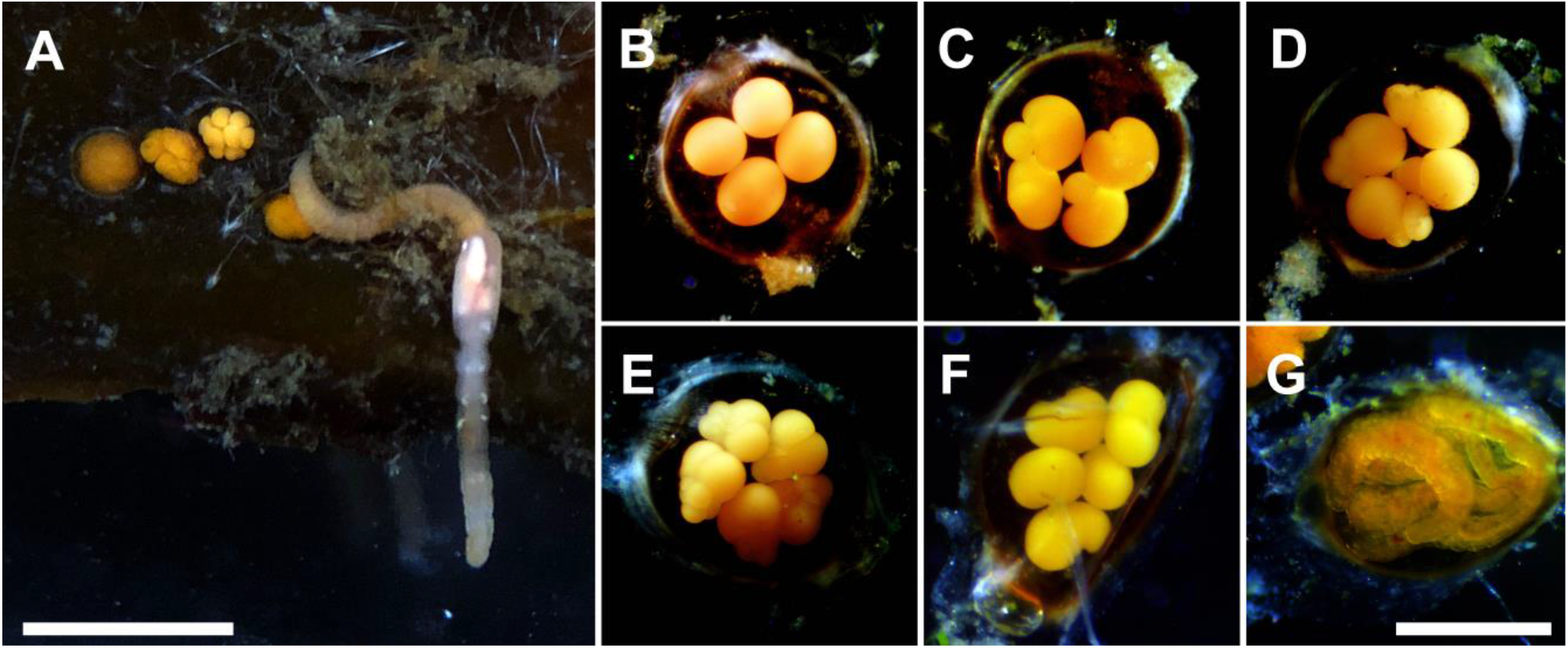
The White Sea littoral oligochaete *Lumbricillus* sp. and its normal development. (A) An adult worm on the surface of a brown algae thallus, along with several clutches of eggs. Scale bar: 5 mm. (B-G) Cocoons with embryos at different stages of development – zygote (B), 2-cell (C), 4-cell (D), 8-cell (E), gastrula (F), segmented juvenile (G). Scale bar: 1 mm.

A morphological analysis conducted in accordance with the identification key (Chekanovskaya, 1962) revealed that these oligochaetes can be classified within the genus *Lumbricillus*, family Enchytridae. S-shaped unidentate chaetae, soldered nephridia, dorsal blood vessel beginning behind the clitellum, large lobed testes were revealed as determinative characters for this genus. In order to more accurately establish the taxonomic affiliation of the species under study, adult oligochaetes (N=5), as well as their cocoons (N=10), were subjected to molecular phylogenetic analyses using both nuclear and mitochondrial markers. Consequently, it was determined that the DNA samples from the adult worms and cocoons were identical and belonged to the same species. The phylogenetic analysis of two molecular markers indicates that this species forms a distinct clade in both phylogenetic trees. This clade is part of a “core group” within the paraphyletic genus *Lumbricillus*. In the COI-based tree, this species exhibits a sister relationship with an undescribed *Lumbricillus* sp. from the UK, sequence F (Fig. 2A). In the 18S tree, it clusters with *Lumbricillus cf. helholandicus* (Fig. 2B). In both cases, these relationships have low bootstrap support, and the p-distances are significant. Two of the three species of this genus that have been described for the White Sea (Krasnova et al., 2010), *Lumbricillus pagenstecheri* and *Lumbricillus lineatus*, are classified in two different clades on phylogenetic trees. However, they are not closely related to the samples under study. The third species known from the White Sea, *Lumbricillus murmanicus*, is regrettably absent from the GenBank database and also absent from the known European oligochaete identification keys (Chekanovskaya, 1962; van Haaren, Soors, 2013; Schmelz, Collado, 2010). It can thus be concluded that the investigated littoral oligochaete can be classified as representatives of the species *Lumbricillus murmanicus*, which was previously described from the Murman coast, or representatives of another known species not included in the current WSBS fauna lists, or they can be considered to represent a new scientific discovery. At this stage, further identification of this species is not possible, as there have been no integrative taxonomic studies of this genus to date. Consequently, we designate this specimen as *Lumbricillus* sp., which differs from the species currently documented in the databases.

**Figure 2.**
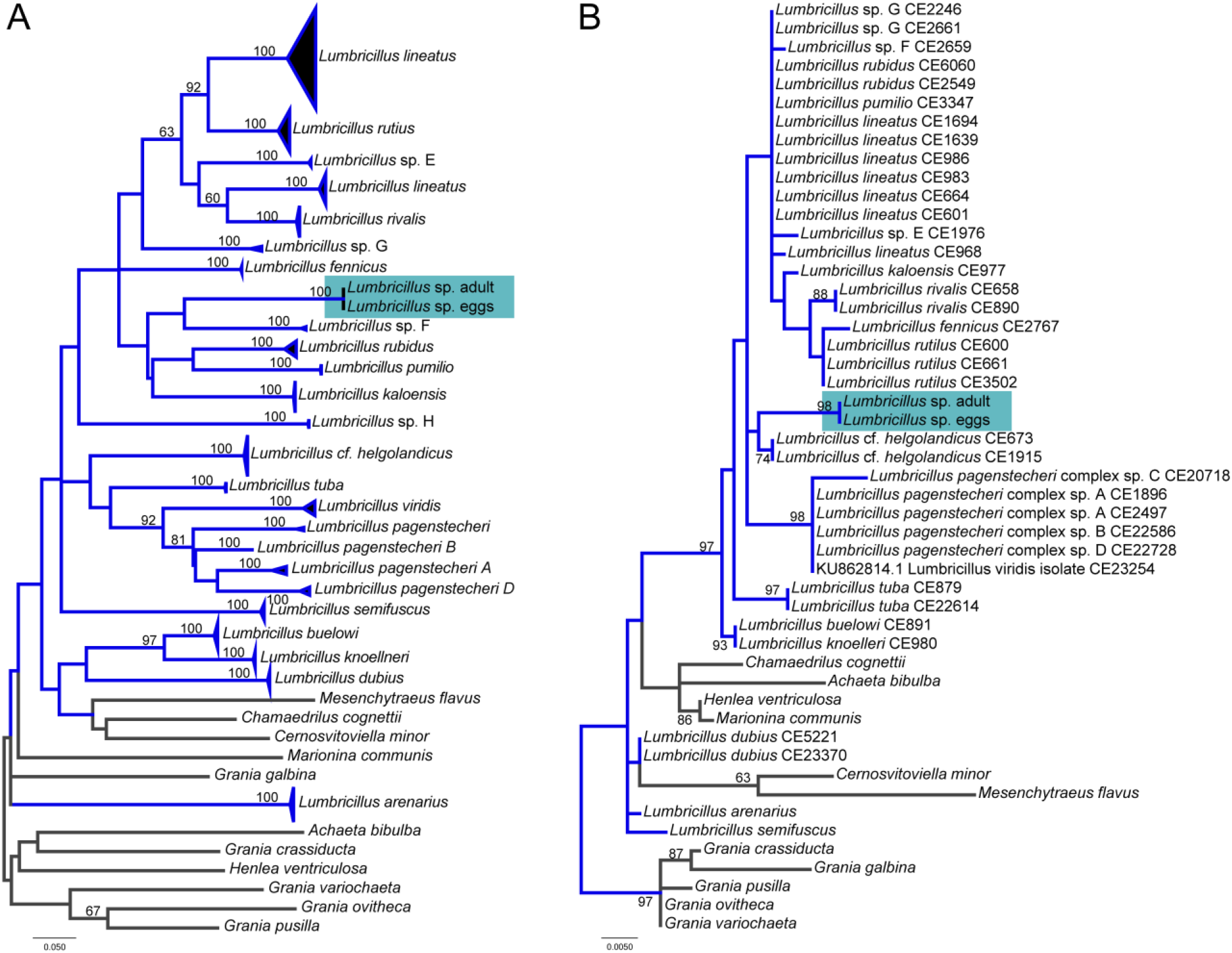
Phylogenetic trees constructed based on the comparison of the sequences of the nuclear 18S rRNA (A) and the mitochondrial COI gene (B). Species-level clades of the genus *Lumbricillus* are collapsed to a single branch except the target taxon on (A). Bootstrap values are shown in respective nodes (only values >60% are shown). Studied specimens are highlighted by a colored box. Blue branches refer to the genus *Lumbricillus*, grey 3 to outgroup taxa.

**Figure 3.**
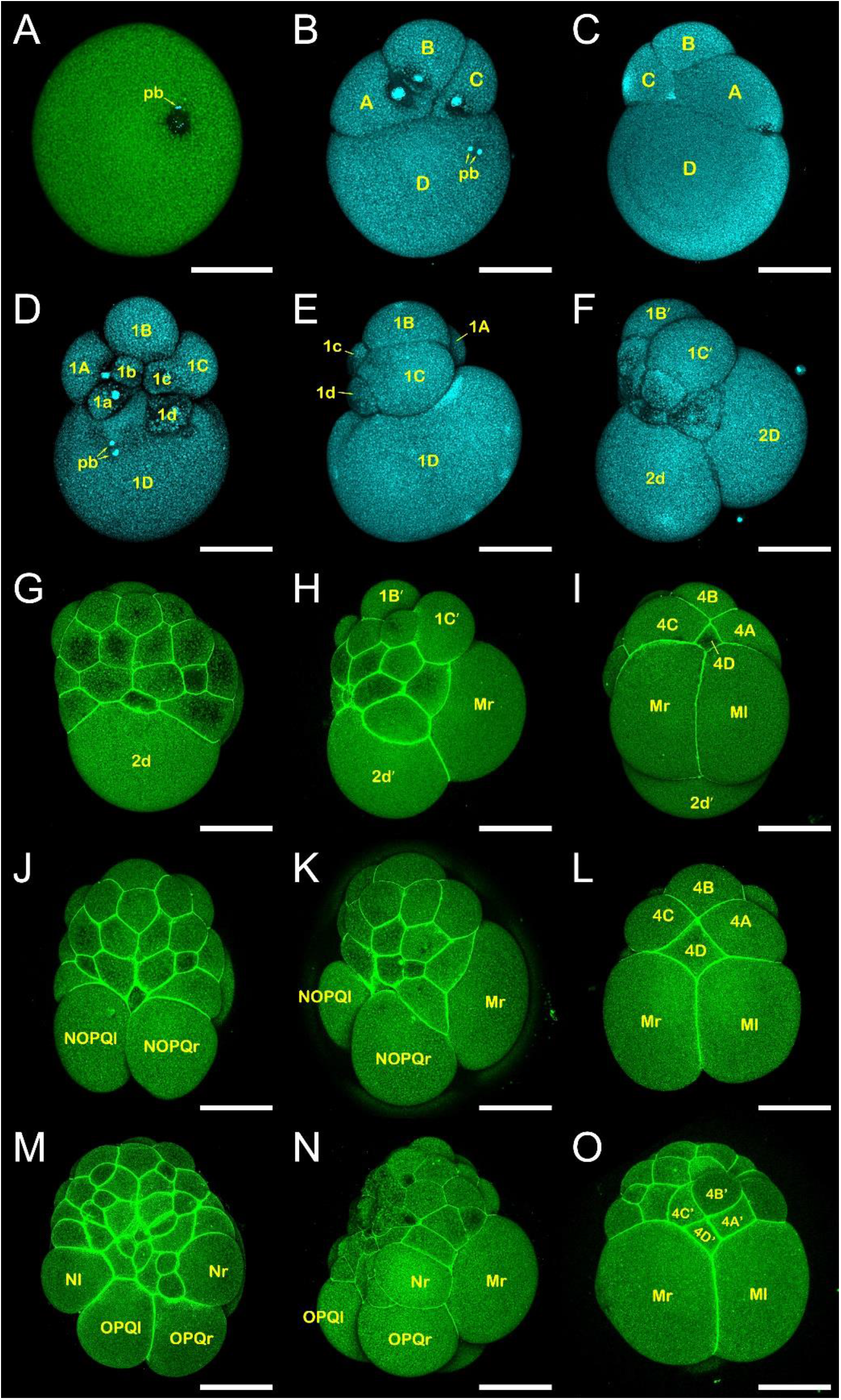
Confocal images of cleavage embryos of *Lumbricillus* sp. (A) Zygote, observation from the animal pole. A distinct region of ooplasm devoid of yolk and containing nuclear material is discernible. (B-C) 4-cell embryo, view from the animal pole (B) and from the vegetal pole (C). The heteroquadrant pattern is clearly visible, with blastomere D exhibiting a significantly larger size than the other blastomeres. (D-E) 8-cell embryo, view from the animal pole (D) and from the vegetal pole (E). The macromeres (1A-1D) and small micromeres on the animal side (1a-1d) are clearly visible. A quartet of micromeres does not form a compact rosette. (F) 9-cell embryo, view from the right side. The division of 1D-cell has resulted in the formation of an equal-sized 2D macromere and 2d micromere. (G-I) The embryos at the stage of mesoteloblasts (Mr and Ml) formation in three views: from the animal pole (G), from the right side (H), and from the vegetal pole (I). (J-L) The embryos at the stage of ectoteloblasts (NOPQl and NOPQr) formation: from the animal pole (J), from the right side (K), and from the vegetal pole (L). (M-O) The embryos at the stage of N-ectoteloblasts formation: from the animal pole (M), from the right side (N), and from the vegetal pole (O). The scale bar is 100 μm. The pseudo-colours indicate the following: green (A, G-O) represents the staining of cell borders with phalloidin, cyan (A-F) represents nuclear staining with DAPI, pb: polar bodies.

The study of embryonic development was conducted on freshly collected material under laboratory culture conditions (filtered seawater, 10°C). Embryos at all stages of development are inherently fragile, rendering them susceptible to trauma upon removal from their cocoons. The dense cocoon shell can cause significant damage to embryos, and the subsequent exposure to seawater results in their rapid deterioration and demise. In this instance, the contents of the destroyed embryonic cells, which occupy the entirety of the space beneath the thin fertilisation membrane, are clearly visible. Therefore, the embryos were not removed from their cocoons in order to track the timing of their development and observe their normal development. The majority of oligochaetes are hermaphroditic, exhibiting cross-fertilisation behaviors. During copulation, eggs and sperm are deposited in a cocoon-like structure formed in the region of the clitellum. As a result, fertilisation occurs and the zygote is formed during cocoon deposition, allowing embryonic development to be observed from its earliest stages. Given this, the approximate time of stage onset was determined in hours post-oviposition (hpo). The maximum time of onset of the two-cell stage (Fig. 1C) was determined to be 6-7 hours after the cocoons were collected, which was taken as the 6-7 hpo. Visual observations revealed that cleavage occurred in an unequal and largely asynchronous manner. The four-cell stage occurs at 12-14 hpo (Fig. 1D), the eight-cell stage at 17-20 hpo (Fig. 1E), and the ∼16-cell stage at 38-40 hpo. During the following days of development, teloblastogenesis and germ band formation occur, resulting in an elongated embryo with the characteristic bean shape (Fig. 1F). The fully formed, segmented young worms emerge from the cocoons and actively crawl away (Fig. 1G).

In order to gain greater insight into the early stages of embryonic development, the morphology of cells at different developmental stages was examined using a laser scanning confocal microscope. The fertilisation membrane was observed to fit the embryos in a tight manner. Neither mechanical nor chemical methods of removing the fertilization shell (treatment with 2.5% L-cysteine, 0.1% pronase or 1% sodium thioglycolate) were effective for them. In order to facilitate microscopic examination, the embryos were stained with low molecular weight fluorescent dyes that were capable of penetrating the fertilisation membrane. Cell borders were stained with fluorescently labelled phalloidin, while nuclei were identified with propidium iodide or DAPI (Fig. 3).

In *Lumbricillus* embryos at the zygote stage and at the first cleavage divisions (Fig. 3A-C), there are no indications of pole plasma formation. In contrast to the distribution of polar plasm in the D blastomere during ooplasmic segregation observed in *Tubifex*, no cellular material, including yolk granules, undergoes an uneven distribution between blastomeres in *Lumbricillus*. Its cleavage, as is characteristic of oligochaetes, exhibits a distinct heteroquadrant spiral pattern. The first cleavage division is markedly unequal, resulting in the formation of a small AB cell and a large CD cell (Fig. 1 C). The CD cell subsequently divides, resulting in the formation of the smaller C cell and the larger blastomere D, which accounts for approximately 80% of the embryo’s volume. In the subsequent phase of the first cleavage, the AB cell divides into A- and B-blastomeres of equal size, leading to the establishment of the four-cell stage (Fig. 3 B-C). During the next round of cleavage, all 4 blastomeres undergo asynchronous unequal divisions, giving rise to the large macromeres (1A, 1B, 1C and 1D) on the vegetal side of the embryo and the small micromeres (1a, 1b, 1c and 1d) on the animal side (Fig. 3 D-E). Soon after, the 1D cell divides into two equal-sized cells, the 2D macromere and the 2d micromere (Fig. 3 F). The micromere 2d is the primary somatoblast – it contacts the animal micromeres of the first quartet and subsequently gives rise to ectoteloblasts. Concurrently, the 2D macromere contacts the major quartet in the vegetal zone and will soon produce mesoteloblasts.

In the subsequent developmental phase, the progeny of the 2d blastomere (2d′) and the main quartet cells (1A′, 1B′, and 1C′) rapidly produce a mass of small ectodermal cells at the animal pole of the embryo (Fig. 3 G-H). A series of highly unequal divisions of the 2D macromere results in the formation of a large 4d micromere and a small 4D macromere, which is in contact with macromeres A, B and C (Fig. 3 I, L). The 4d cell then divides bilaterally into two large mesoteloblasts (Ml and Mr cells) (Fig. 3 I, L). The M cells are situated ventrally on both sides of the embryo, which disrupts the spiral cleavage pattern. Following this, the 2d cells also undergo an equal division, resulting in a bilaterally symmetrical pair of ectoteloblasts precursors (NOPQ cells) (Fig. 3 J-K, M-O). Each of these will subsequently give rise to four ectoteloblasts, N-, O-, P-, Q-cells (Fig. 4 A-B), which will start to form primary blast cells columns (or bandlets). The aggregated bandlets from N, O, P and Q teloblasts form the ectodermal germ band and M cells form the mesodermal germ band which underlines the ectodermal one (Fig. 4). The gastrulation includes germ bands elongating, then curving toward the ventral midline, and closing from anterior to posterior end on the ventral midline along with body elongation (Takahashi et al., 2008).

**Figure 4.**
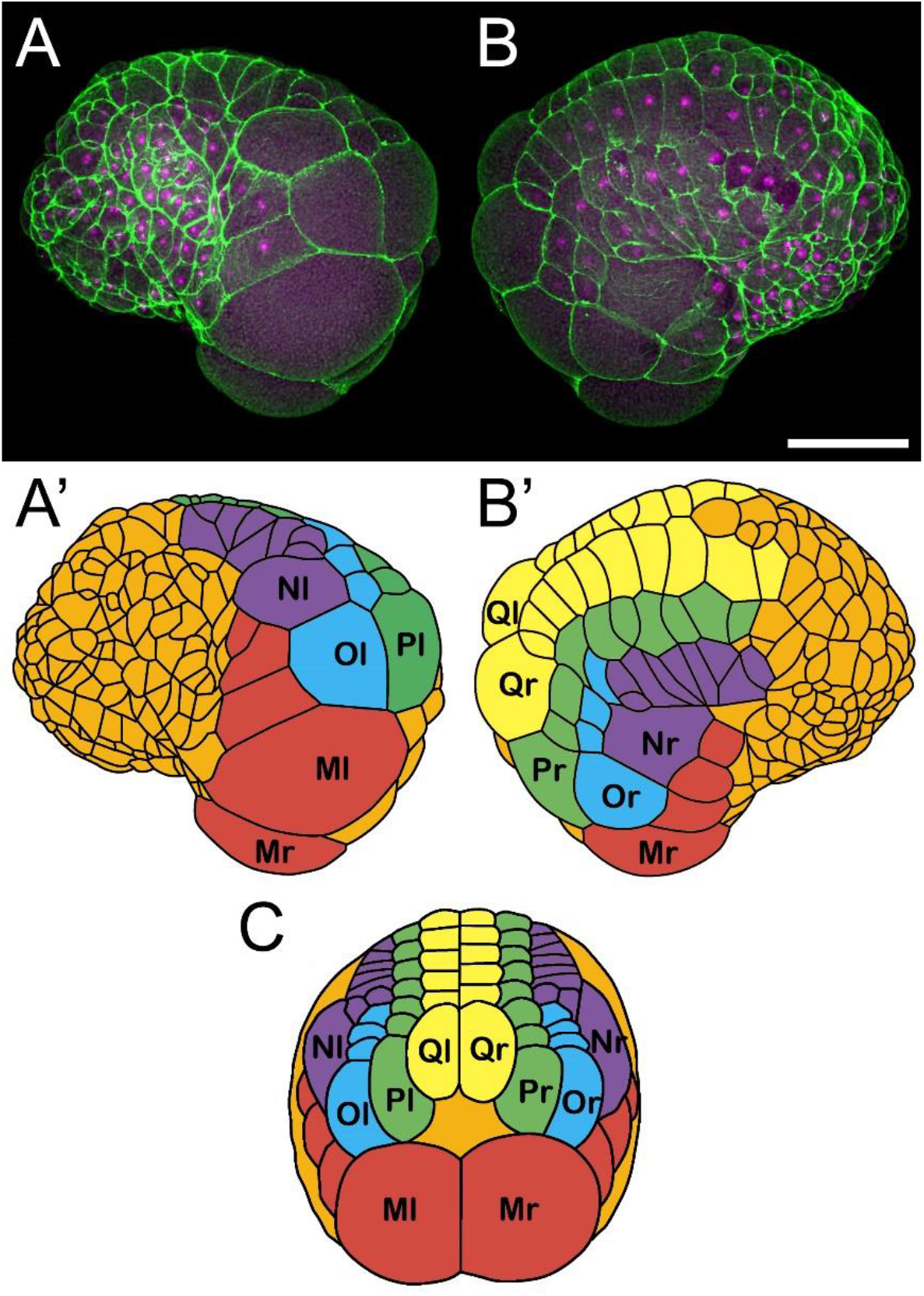
Germ bands formation in embryo of *Lumbricillus* sp. (A-B) Confocal images and their graphical drawings (A’-B’) showing the same embryo, view from the left (A) and right (B) side. (C) A schematic representation of the mutual arrangement of teloblasts at the posterior end of the embryo at the specified stage of development. The scale bar is 100 μm. The pseudo-colours indicate the following: green represents the staining of cell borders with phalloidin and magenta represents nuclear staining with propidium iodide.

Among oligochaetes, embryonic development has been most comprehensively described for two model species: *Tubifex tubifex* and *Enchytraeus coronatus* (Aoki, Shimizu, 2017; Yoshida et al., 2019). The embryonic development of *Lumbricillus* sp. displays oligochaete-specific traits but is more consistent with the described development of *Enchytraeus* (Berger et al., 2004). In particular, no evidence was found of pole plasma, no rosette formation by the four micromeres of the first quartet (Fig. 3 D), equal size of the 2d and 2D blastomeres (Fig. 3 F), and a small size of the 4D blastomere in comparison to the large 4d blastomere. In contrast to the high developmental variability observed in *Tubifex* (Meshcheryakov, 1990), the embryos of *Lumbricillus* develop in a highly stable and synchronous manner. All embryos within a single cocoon are at the same developmental stage, exhibiting an almost identical pattern of cell arrangement (Fig. 1 D-E). The combination of synchrony and low developmental variability, coupled with the accessibility of the object, ease of maintenance and manipulation, makes *Lumbricillus* sp. a highly promising subject for the study of embryonic development in Annelides.

## CONCLUSIONS

This study presents the initial description of the early development of the littoral oligochaete, identified as *Lumbricillus* sp., through the utilisation of in vivo light microscopy observation and confocal microscopy. The data obtained here constitute an essential prerequisite for further studies of this species. The embryonic development of *Lumbricillus* sp. exhibits numerous features characteristic of “classical” oligochaete development, including unequal heteroquadrant cleavage and the early establishment of the axial organization of the embryo, as well as the formation of meso- and ectoteloblasts and germ bands. Concurrently *Lumbricillus* sp. exhibits specific characteristics that are typical of other representatives of the Enchytriidae family, such as equal size of the 2d and 2D blastomeres, and a small size of the 4D blastomere in comparison to the large 4d blastomere. Futhermore, this oligochaete exhibits highly stable and synchronous development, and requires relatively simple conditions for the obtaining and maintenance of embryonic material. This makes *Lumbricillus* sp. a promising object for developmental, ecological and evolutionary studies amongst marine representatives of Clitellata that can make a significant contribution to the comparative study of the biology of the Annelida group.

## CONTRIBUTIONS

E.P.M. – investigation (sample collection, embryonic culture, developmental timing, microscopic observation), data curation, writing - original draft preparation. T.V.N. – investigation (DNA sequencing), resources, funding acquisition. I.A.E. – investigation (phylogenetic analysis), software, validation, data curation, visualization. V.V.K. – investigation (fluorescent microscopy), visualization. M.L.S. – supervision, resources. D.A.N. - conceptualization, writing - review and editing, project administration. All authors have read and agreed to the published version of the manuscript.

## ACKNOWLEDGEMENTS

The light microscopy studies were conducted using equipment of the Center of microscopy WSBS MSU. The authors are deeply grateful to the White Sea Biological Station for providing their infrastructure and equipment for collecting and processing the samples.

## ETHICS DECLARATIONS

### Ethics Approval And Consent To Participate

The researchers followed the norms for the use of animals in experiments adopted at the Koltzov Institute of Developmental Biology of the Russian Academy of Sciences, which are regulated by the Institutional Bioethics Committee and comply with international standards, in particular ASPA 1986 (Animals (Scientific Procedures) Act 1986, https://www.legislation.gov.uk/ukpga/ 1986/14/contents).

